# TRPV4 activation enhances LPS-induced IL10 production to suppress excessive microglial activation

**DOI:** 10.1101/2025.05.23.655698

**Authors:** Kosei Tamada, Eri Tomisawa, Saori Kuwata, Takahito Miyake, Kazuki Nagayasu, Shuji Kaneko, Koji Shibasaki, Hisashi Shirakawa

## Abstract

Microglia are resident immune cells in the brain. Under pathological conditions, activated microglia trigger neurodegeneration by secreting pro-inflammatory molecules such as NO, TNFα, IL6 and IL1β. In contrast, microglia also possess a negative feedback mechanism to prevent the excessive inflammation by secreting anti-inflammatory cytokines such as IL10. It is reported that activation of transient receptor potential vanilloid 4 (TRPV4), one of the non-selective cation channels, suppresses the lipopolysaccharide (LPS)-induced microglial activation. However, the effects of TRPV4 activation on the microglial anti-inflammatory responses remain unknown. In this study, we investigated whether TRPV4 contributes microglial anti-inflammatory responses using primary cultured murine microglia. We found that application of a selective TRPV4 agonist, GSK1016790A, enhanced LPS-induced IL10 release in the microglia, which was fully suppressed by co-application of a selective TRPV4 antagonist, GSK2193874, or absent in cultured microglia derived from TRPV4-knockout (TRPV4-KO) mice. Furthermore, neutralization of extracellular IL10 significantly reversed suppressive effects of GSK1016790A on LPS-induced release of TNFα, IL6 and IL1β. Expression pattern of microglial activation markers implied that GSK1016790A shifted microglial activation status to an immunoregulatory phenotype. Taken together, these results indicate that microglial TRPV4 plays an important role in promoting the production of LPS-induced IL10, which results in suppressing excessive inflammation in an autocrine or paracrine fashion.

## 1. Introduction

Microglia, the resident immune cells of the central nervous system (CNS), play an important role in maintaining brain homeostasis by monitoring neuronal function, and clearing apoptotic cells and debris in the healthy brain [1]. Microglia are activated in response to environmental cues and produce several cytokines [2,3]. Activated microglia which produce pro-inflammatory cytokines such as TNFα, IL6 and IL1β defend the brain against pathogens and tumor cells [2,4,5]. On the other hand, there are also microglia which produce anti-inflammatory cytokines such as IL10, TGFβ and IL4, which have an important role in not only promoting tissue repair and angiogenesis but also providing a negative feedback mechanism [2,5–8]. Especially, IL10 is known as a key immunoregulatory cytokine that mediates a negative feedback loop [8,9], and an insufficiency of IL10 production or signaling followed by ongoing excessive activation of microglia which produce pro-inflammatory cytokines is a common feature of the pathogenesis and pathology of various neurodegenerative diseases including Parkinson’s disease [10–12], multiple sclerosis [13–15] and neuropathic pain [16,17]. In this context, to understand how microglia regulate its negative feedback mechanism during inflammation is critical for the development of novel immune modulatory strategies.

Transient receptor potential vanilloid 4 (TRPV4) channel is a nonselective cation channel that is activated by various physical and chemical stimuli, including hypoosmolarity, warm temperature, mechanical stimulation, and arachidonic metabolites [18,19]. In the brain, microglia show the highest TRPV4 mRNA expression [19,20]. It is previously reported that activation of microglial TRPV4 by 4α-phorbol 12,13-didecanoate (4α-PDD), a relatively poor selective TRPV4 agonist, suppresses the LPS-induced activation by depolarization of the resting potential, leading to the attenuation of the driving force for Ca^2+^ influx [20]. However, the molecular details how TRPV4 activation suppresses the microglial activation are still elusive. Especially, the effects of TRPV4 activation on the microglial anti-inflammatory responses have not been addressed yet.

In this study, we hypothesize that activation of TRPV4 augments the production of anti-inflammatory cytokines, which is followed by the suppression of microglial activation. Using primary cultured microglia derived from wild-type (WT) or TRPV4-knockout (TRPV4-KO) mice, we demonstrate that microglia treated with LPS and GSK1016790A, a highly selective TRPV4 agonist, show a unique type of activation by producing large amounts of IL10, which may contribute to the suppression of the LPS-induced pro-inflammatory cytokine production in microglia.

## 2. Results

### 2.1. TRPV4 is functionally expressed in microglia

First of all, we investigated whether primary cultured murine microglia possess TRPV4. TRPV4 mRNA was expressed in microglia (Figure 1A). Fura-2-based Ca^2+^ imaging experiments revealed that application of a selective TRPV4 agonist GSK1016790A (1 μM) induced an increase of intracellular Ca^2+^ concentration ([Ca^2+^]_i_) in WT microglia (Figure 1B), which was almost completely inhibited by a selective TRPV4 antagonist GSK2193874 (1 μM) (Figure 1C). By contrast, the GSK1016790A-induced [Ca^2+^]_i_ increase was not observed in TRPV4-KO microglia (Figure 1D). Whole-cell patch clamp recordings showed that GSK1016790A (0.1 μM) evoked outwardly rectified currents (Figure 1E), which were not observed in TRPV4-KO microglia (Figure 1F). These results indicate that TRPV4 is located and functional on the plasma membrane of microglia.

**Figure 1.**
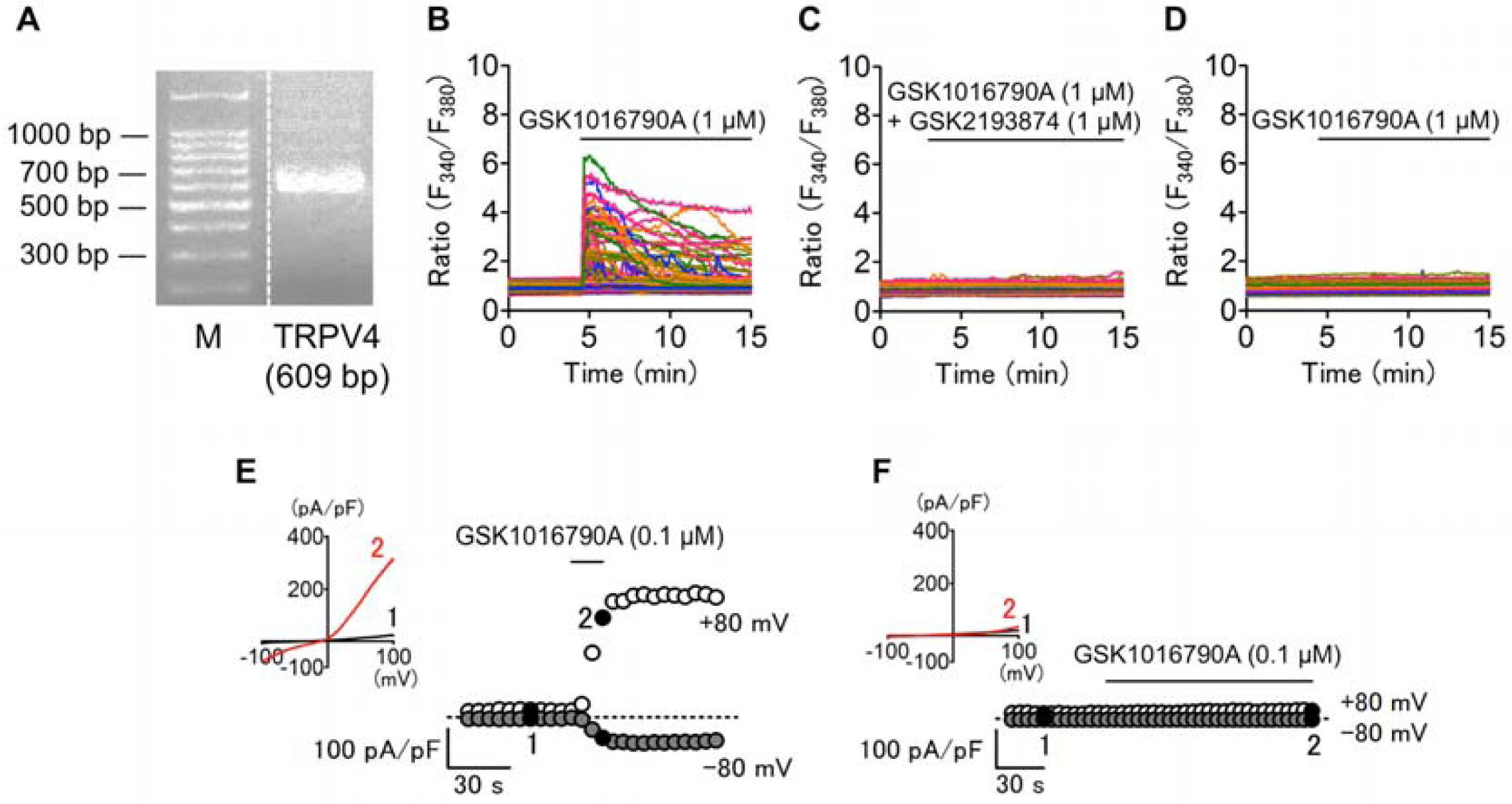
Functional expression of TRPV4 in cultured murine microglia. (A) Representative result of RT-PCR using cultured murine microglia. Marker (M) shows 100 bp DNA size ladder. (B-D) Representative traces of intracellular Ca^2+^ imaging experiments from WT (B, C) or TRPV4-KO (D) microglia. (E, F) Representative whole-cell currents from WT (E) or TRPV4-KO (F) microglia. Membrane potential was 0 mV. Insets in E and F indicate current–voltage relationships at the time indicated by black-filled circles (1: before, 2: after application of GSK1016790A). GSK1016790A, a selective TRPV4 agonist; GSK2193874, a selective TRPV4 antagonist. Drugs were applied during the periods indicated by the horizontal lines.

### 2.2. TRPV4 activation increases the LPS-induced IL10 mRNA expression in microglia

To clarify the role of TRPV4 in the negative feedback mechanisms in microglia, we investigated whether the activation of TRPV4 affects the expression levels of various anti-inflammatory cytokines. In quantitative RT-PCR analysis, we found that only IL10 mRNA was significantly increased in WT microglia cotreated with LPS (100 ng/mL) and GSK1016790A (1 μM), compared with WT microglia treated with LPS (Figure 2A), which was not observed in TRPV4-KO microglia (Figure 2B). mRNA levels of other anti-inflammatory cytokines such as TGFβ and IL4 did not change (Figure 2C, D). These results suggest that TRPV4 activation specifically promotes IL10 mRNA expression in the LPS-stimulated microglia.

**Figure 2.**
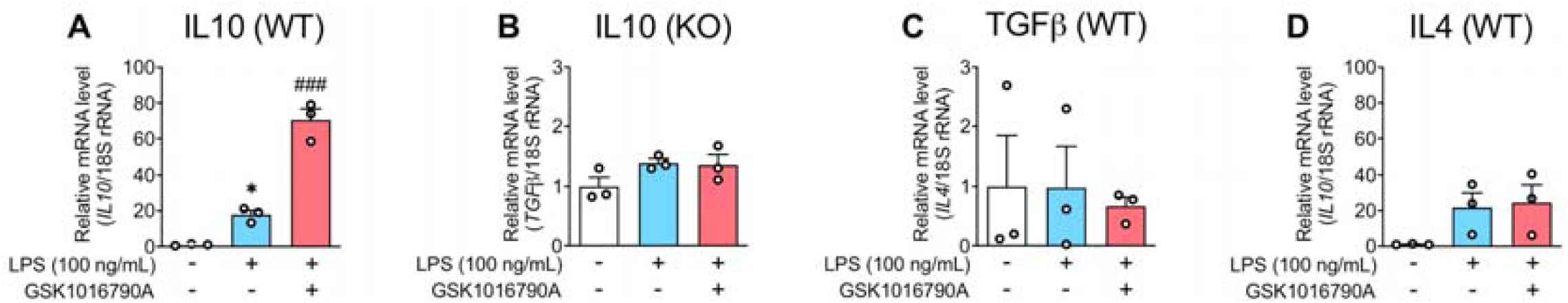
Effects of TRPV4 activation on the mRNA expression levels of anti-inflammatory cytokines in microglia. WT (A,C,D) or TRPV4-KO (KO,B) microglia were treated with or without GSK1016790A (1 μM) in addition to LPS (100 ng/mL) for 24 h. The relative mRNA levels of IL10 (A,B), TGFβ (C) and IL4 (D) were determined by quantitative RT-PCR. *P < 0.05 vs non-treated, ^###^P < 0.001 vs LPS alone, n = 3. Data are presented as means ± SEM.

### 2.3. TRPV4 activation augments the LPS-induced IL10 release in microglia

Next, we examined the effect of GSK1016790A on the LPS-induced IL10 release in microglia, and found that GSK1016790A increased the LPS-induced IL10 release in a concentration-dependent manner (Figure 3A). The augmentation of the LPS-induced IL10 release by GSK1016790A (1 μM) was fully inhibited by cotreatment of GSK2193874 (1 μM). In addition, GSK2193874 by itself also suppressed the LPS-induced IL10 release (Figure 3B). On the other hand, the level of LPS-induced IL10 in TRPV4-KO microglia was comparable to that of WT microglia, but GSK1016790A had no effects (Figure 3C). Of note, GSK1016790A had no effect on the cell viability of WT microglia (Figure 3D). These results indicate that TRPV4 activation promotes the LPS-induced IL10 release in microglia.

**Figure 3.**
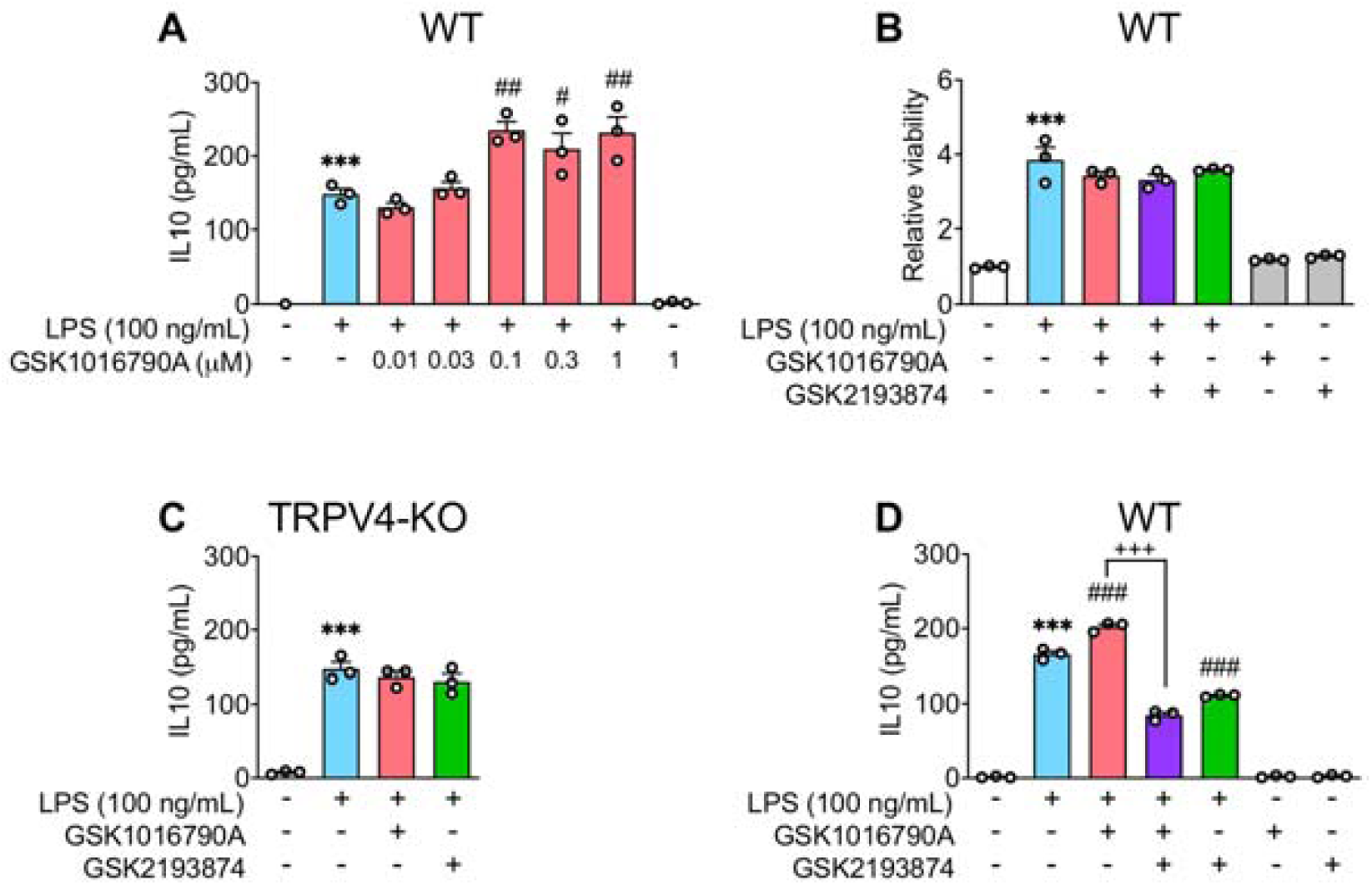
TPRV4-dependent modulation of the LPS-induced IL10 release in microglia. (A) WT microglia were treated at indicated concentrations of GSK1016790A in addition to LPS (100 ng/mL) for 72 h. (B, C) Effects of 72 h application of GSK1016790A (1 μM) and/or GSK2193874 (1 μM) on the LPS (100 ng/mL)-induced IL10 release in WT (B) or TRPV4-KO (C) microglia. (D) Effects of 72 h application of GSK1016790A (1 μM) and/or GSK2193874 (1 μM) with or without LPS (100 ng/mL) on cell viability. The concentrations of IL-10 in the supernatants were determined by ELISA. The relative cell viability was determined by MTT assay. ***P < 0.001 vs non-treated, ^#^P < 0.05, ^##^P < 0.01, ^###^P < 0.001 vs LPS alone, ^+++^P < 0.001, n = 3. Data are presented as means ± SEM.

### 2.4. TRPV4-dependent IL10 suppresses the LPS-induced pro-inflammatory cytokine release in microglia

We explored whether endogenous IL10 released after TRPV4 activation contributes to the TRPV4-dependent suppressive effect on the release of pro-inflammatory cytokines such as TNFα [15]. The LPS-induced release of IL6 and IL1β as well as TNFα was significantly suppressed in the presence of GSK1016790A (Figure 4A-C). The inhibitory effects of GSK1016790A on the release of these cytokines were fully attenuated by cotreatment with GSK2193874. We further examined the effects of endogenous IL10 on the release of these cytokines. Neutralization of IL10 by a specific antibody against IL10 (αIL10 IgG, 50 ng/ml) significantly blocked the inhibitory effects of GSK1016790A on TNFα, IL6 and IL1β, while an isotype control antibody (control IgG, 50 ng/ml) had no effect (Figure 4D-F). These results indicate that the enhanced release of IL10 after TRPV4 activation may contribute to the suppression of pro-inflammatory cytokine release.

**Figure 4.**
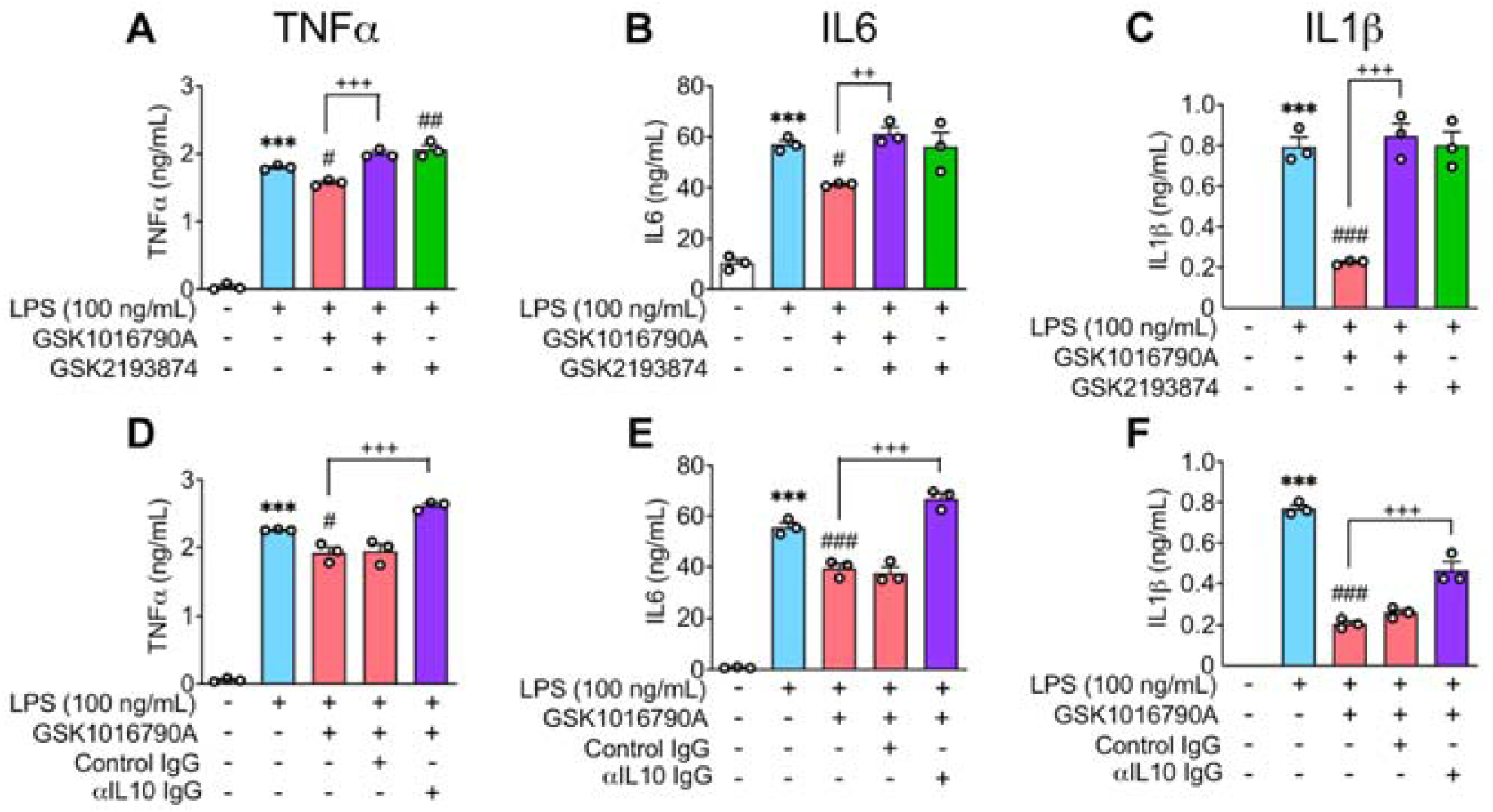
Effects of TRPV4 activation and neutralizing IL10 on the LPS-induced pro-inflammatory cytokine release in microglia. (A−C) WT microglia were treated with GSK1016790A (1 µM) and/or GSK2193874 (1 μM) in addition to LPS (100 ng/mL) for 72 h. (D-F) WT microglia were treated with GSK1016790A (1 μM) with or without a neutralizing antibody against IL10 (αIL10 IgG, 50 ng/mL) or its isotype control antibody (50 ng/mL) in addition to LPS (100 ng/mL) for 72 h. The concentrations of TNFα (A, D), IL6 (B, E) and IL1β (C, F) in the supernatants were determined by ELISA. ***P < 0.001 vs non-treated, ^#^P < 0.05, ^##^P < 0.01, ^###^P < 0.001 vs LPS alone, ^++^P < 0.01, ^+++^P < 0.001, n = 3. Data are presented as means ± SEM.

### 2.5. TRPV4-dependent IL10 suppresses the LPS-induced pro-inflammatory cytokine release in microglia

We then sought to elucidate the molecular mechanisms underlying the TRPV4-dependent microglial IL10 release especially focusing on the role of Ca^2+^, since Ca2+ is a well-known molecule to regulate the function of microglia [21–23]. However, an intracellular Ca^2+^ chelator BAPTA-AM (1-3 μM) had no effect on the LPS-induced and GSK1016790A-potentiated release of IL10. In addition, BAPTA-AM did not affect the LPS-induced IL10 release by itself (Figure 5). Moreover, a Ca^2+^ ionophore ionomycin (1-3 μM) significantly suppressed the LPS-induced IL10 release in a concentration-dependent manner (Figure 5). These results suggest that, although Ca^2+^ in itself has an inhibitory effect on the release of IL10, the effects of Ca^2+^ influx following TRPV4 activation may be negligible in the TRPV4-dependent IL10 release in microglia, consistent with a previous study [20].

**Figure 5.**
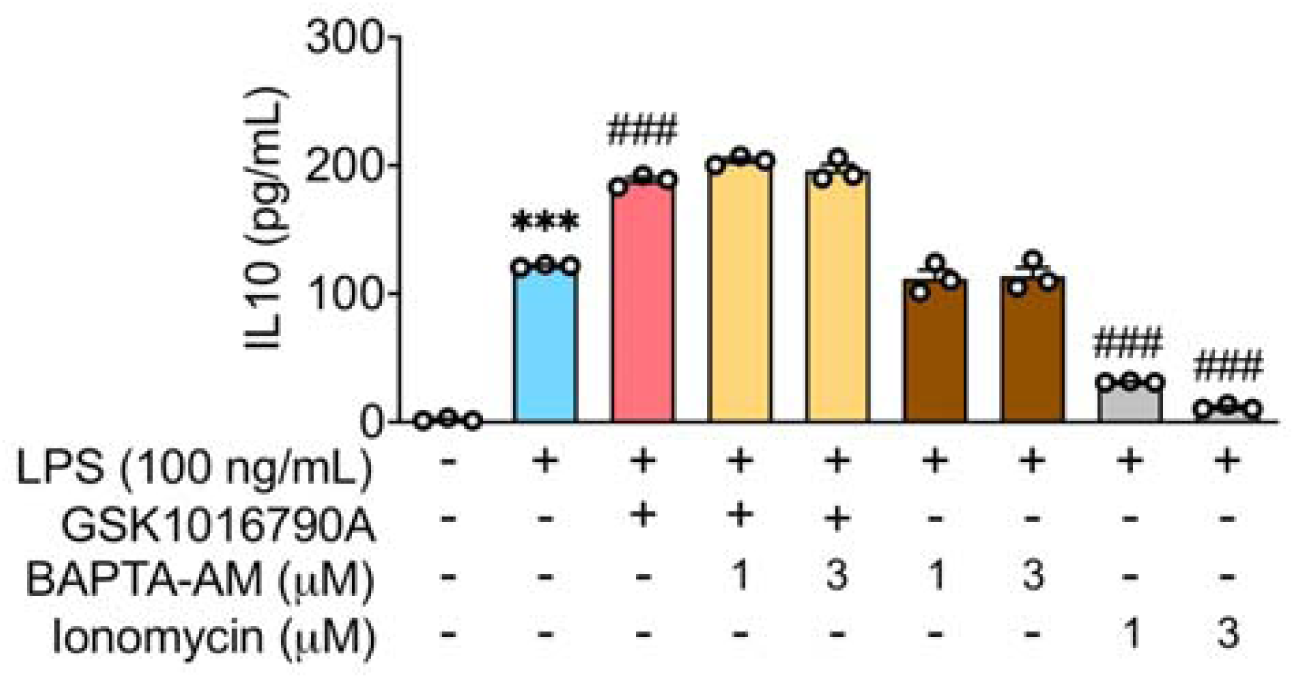
Role of Ca^2+^ ions in the LPS-induced IL10 release in microglia. WT microglia were treated with indicated concentrations of an intracellular Ca^2+^ chelator BAPTA-AM or a Ca^2+^ ionophore ionomycin with or without GSK1016790A (1 μM) in addition to LPS (100 ng/mL) for 72 h. The concentrations of IL-10 in the supernatants were determined by ELISA. ***P < 0.001 vs non-treated, ^###^P < 0.001 vs LPS alone, n = 3. Data are presented as means ± SEM.

### 2.6. TRPV4 activation induces the expression of regulatory microglial markers

Finally, we examined the effects of GSK1016790A on the expression of pro/anti-inflammatory markers in microglia. We checked a major marker of pro-inflammatory microglia, CD86, and found that CD86 mRNA was significantly increased in LPS- and GSK1016790A-cotreated WT microglia, compared with LPS-treated WT microglia by quantitative RT-PCR analysis (Figure 6A). On the other hand, a major marker of anti-inflammatory microglia, CD206 was not changed at mRNA level (Figure 6B). It is known that some IL10^high^ microglia do not express CD206 but CD86 [24,25]. Therefore, we examined sphingosine kinase-1 (SPHK1), which has been reported as a marker of IL10^high^ CD86^+^ microglia [26–28]. SPHK1 mRNA was not significantly but almost doubled in LPS and GSK1016790A-treated WT microglia compared with LPS-treated WT microglia (Figure 6C). These results suggest that TRPV4 activation may be associated with changes to IL10^high^ CD86^+^ microglia.

**Figure 6.**
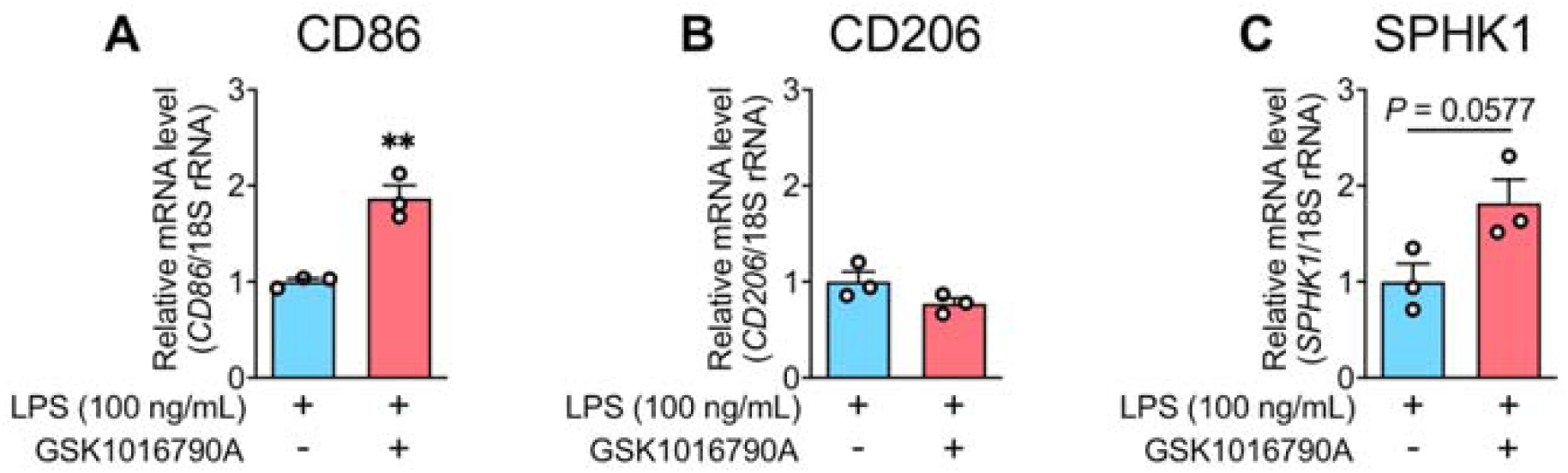
Effects of TRPV4 activation on the expression of microglial polarization markers. WT microglia were treated with or without GSK1016790A (1 μM) in addition to LPS (100 ng/mL) for 24 h. The relative mRNA levels of CD86 (A), CD206 (B) and SPHK1 (C) were determined by quantitative RT-PCR. **P < 0.01 vs LPS alone, n = 3. Data are presented as means ± SEM.

## 3. Discussion

In this study, we suggest that TRPV4 activation results in a shift in microglial activation state to IL10^high^ CD86^+^ leading to the enhancement of LPS-induced IL10 production in microglia. The TRPV4-dependent IL10 production may have an important role in the inhibition of LPS-induced pro-inflammatory cytokine production.

Although we used a synthetic agonist GSK1016790A to open TRPV4 channels, the endogenous ligands for microglial TRPV4 are obscure. TRPV4 is known to be directly activated by warm temperature [29] or LPS itself [30]. In this study, GSK2193874 suppressed the LPS-induced IL10 release at 37°C, but there was no difference between WT and TRPV4-KO microglia in the LPS-induced IL10 release, indicating that both warm temperature and LPS may contribute to TRPV4 activation in microglia, and that TRPV4-KO microglia may possess some alternative mechanisms to produce IL10 in response to LPS at 37°C. However, endogenous ligands other than warm temperature and LPS should exist, since GSK1016790A robustly increased the IL10 release in the presence of LPS at 37°C. In the brain, astrocytic TRPV4 is reported to be activated by arachidonic acid generated at postsynaptic sites [31,32]. As for the production of arachidonic acid, another report showed that activated microglia can release arachidonic acid derivative [33]. It has also been reported that arachidonic acid promotes microglial dysfunction [34]. Thus, it is possible that microglial TRPV4 is activated by arachidonic acid or its derivative from synapses or microglia itself in response to increased lipid metabolism.

Konno *et al.* previously reported that a relatively poor selective TRPV4 agonist 4α-PDD inhibits the LPS-induced production of TNFα from microglia by depolarization of the resting potential, leading to the attenuation of the driving force for Ca^2+^ influx [20]. In this study, we showed that a highly selective TRPV4 agonist GSK1016790A also decreased the LPS-induced IL6 and IL1β release as well as TNFα release in microglia. Furthermore, the neutralization of extracellular IL10 significantly attenuated the effects of GSK1016790A on the TNFα, IL6 and IL1β release. Therefore, the suppressive effects of GSK1016790A on various pro-inflammatory cytokine production may be derived from the IL10 production augmented by TRPV4 activation. We also showed that BAPTA-AM failed to inhibit the effect of GSK1016790A on the LPS-induced IL10 release, and that ionomycin significantly suppressed the LPS-induced IL10 release, indicating that the TRPV4-dependent IL10 production in microglia is independent of Ca^2+^ influx, consistently with Konno *et al.* [20].

Although TRPV4 activation increased the expression of IL10, one of the anti-inflammatory cytokines produced by microglia, it augmented the expression of CD86 but not CD206. This may be explained by the fact that IL10^high^ microglia do not express CD206 but express CD86 [28,35]. These microglia have a central role in producing high levels of anti-inflammatory IL10 [24,28,35–37]. SPHK1 has been proposed as a marker for IL10^high^ CD86^+^ microglia [26–28], and TRPV4 activation increased SPHK1 mRNA expression. These observations suggest that LPS and GSK1016790A-treated microglia exhibit IL10^high^ CD86^+^ phenotype. However, microglia activation of this phenotype is the least understood type, and further investigation is required to clarify the contribution of TRPV4 in leading IL10^high^ CD86^+^ microglia.

In conclusion, this study demonstrated that activation of TRPV4 enhances the LPS-induced IL10 production in microglia presumably by switching the activation state of microglia into IL10^high^ CD86^+^ state. Moreover, the TRPV4-dependent IL10 may have a pivotal role in regulating the production of the LPS-induced pro-inflammatory cytokines. IL10 is known as a key immunoregulatory cytokine that not only mediates a negative feedback loop but also plays several other roles in direct neuroprotective properties [38,39] and mediating effects of immune crosstalk between glial cells [40]. It is reported that IL10 expression in microglia is contributed in the suppression of neurodegeneration in ALS [41], Parkinson’s disease [12], and Alzheimer’s disease [42]. Furthermore, it has been reported that TRPV4 inhibition suppressed pain-like behavior in a mouse model of neuropathic pain [43]. Thus, TRPV4 may be a potential therapeutic target to neurodegenerative diseases accompanied by microglia-related brain inflammation.

## 4. Materials and Methods

### 4.1. Primary cultured microglia

All animals used in this study were treated in accordance with the guidelines of the Kyoto University Animal Experimentation Committee. Microglial cultures were prepared from newborn wild-type or TRPV4 knockout C57BL/6 mice (1–2 days old). Dissociated cells were seeded on 75 cm2 flasks in Dulbecco’s modified Eagle medium (DMEM; D5796, Sigma-Aldrich) supplemented with 10% heat-inactivated fetal bovine serum (Sigma-Aldrich), 5 mg/mL insulin (Biological Industries), and 1% penicillin/streptomycin mixed solution (Nacalai Tesque). The cultures were maintained at 37°C in a humidified 5% CO_2_ atmosphere. After 2–3 weeks, the primary mixed glial cultures were shaken at 150 rpm for 90 min at 37°C. The detached cells were then plated elsewhere at a density of approximately 1.0 × 10^5^ cells/cm^2^ except for 10-mm glass coverslips (4.0 × 10^4^ cells/glass).

### 4.2. Reverse transcription (RT)-PCR

Total RNA was extracted with NucleoSpin RNA (TaKaRa Bio). First-strand cDNA was prepared from total RNA using ReverTra Ace qPCR RT Kit (Toyobo). The cDNA was amplified by PCR using Blend Taq (Toyobo). The set of primers for TRPV4 was 5’-GTTGGAGTCCACCCTGTACG-3’ and 5’-GGCAGCTCCCCAAAGTAGAA-3’.

PCR conditions were as follows: 94°C for 1 min followed by 35 cycles of 94°C for 40 s, 60°C for 40 s and 72°C for 1 min. Amplified PCR products were electrophoresed in 2% agarose gel and visualized under UV light with 0.1 μg/mL ethidium bromide.

### 4.3. Measurement of [Ca^2+^]_i_

Microglia on coverslips were loaded for 40 min with 5 μM fura-2 acetoxymethyl ester (Fura-2 AM; Dojindo) in Krebs-Ringer buffer (140 mM NaCl, 5 mM KCl, 1 mM MgCl2, 2 mM CaCl2, 10 mM glucose and 10 mM HEPES, adjusted to pH 7.4 with NaOH) containing 0.005% cremophore EL (Sigma-Aldrich). The fluorescence images were captured every 5 sec using alternating excitation at 340 and 380 nm and emission at 510 nm with the AQUACOSMOS/ORCA-AG imaging system (Hamamatsu Photonics). All experiments were conducted at room temperature. The ratio of the fluorescent intensity obtained by the excitation/emission of 340 nm/510 nm (F340) to the fluorescent intensity obtained by the excitation/emission of 380 nm/510 nm (F380), namely F340/F380, was calculated to evaluate the intracellular Ca^2+^ concentration.

### 4.4. Whole-cell patch clamp recording

Whole-cell currents were recorded at room temperature with a pipette made from a 1.5-mm outer-diameter glass capillary with a filament (Narishige) and pulled using a P-87 micropipette puller (Sutter). The access resistance ranged from 2 to 5 MΩ when the pipette was filled with the intracellular solution described below. The pipette solution contained 140 mM KCl, 5 mM EGTA, and 10 mM HEPES (adjusted to pH 7.4 with KOH) and the bath solution contained 140 mM NaCl, 5 mM KCl, 2 mM MgCl2, 2 mM CaCl2, 10 mM glucose and 10 mM HEPES (adjusted to pH 7.4 with NaOH). Current-voltage relationships were measured using voltage ramps (−100 to +100 mV over 200 ms, 0.2 Hz). The holding potential was set at 0 mV. Access resistances were compensated up to 70%. Patch-clamp recordings were performed using an EPC-10 patch-clamp amplifier (HEKA Instruments) and PATCHMASTER software (HEKA Instruments). For analysis, data were filtered at 2.9 kHz.

### 4.5. Quantitative RT-PCR

After reverse transcription of total mRNA to cDNA using ReverTra Ace qPCR RT Kit, real-time quantitative PCR was performed using the StepOne real-time PCR System (Life Technologies) and the THUNDERBIRD SYBR qPCR Mix (Toyobo). The sets of primers used for each gene were as follows: 5’-GCAATTATTCCCCATGAACG-3’ and 5’-GGCCTCACTAAACCATCCAA-3’ for 18S ribosomal RNA (rRNA), 5’-GCTGCCTGCTCTTACTGACT-3’ and 5’-CTGGGAAGTGGGTGCAGTTA-3’ for IL10, 5’-TGACGTCACTGGAGTTGTACGG-3’ and 5’-GGTTCATGTCATGGATGGTGC-3’ for TGFβ, 5’-TCTCGAATGTACCAGGAGCCATATC-3’ and 5’-AGCACCTTGGAAGCCCTACAGA-3’ for IL4, 5’-TTGTGTGTGTTCTGGAAACGGAG-3’ and 5’-AACTTAGAGGCTGTGTTGCTGGG-3’ for CD86, 5’-TCTTTGCCTTTCCCAGTCTCC-3’ and 5’-TGACACCCAGCGGAATTTC-3’ for CD206 and 5’-TCCAGAAACCCCTGTGTAGC-3’ and 5’-CAGCAGTGTGCAGTTGATGA-3’ for SPHK1. The temperature cycles were 95°C for 10 min, followed by 40 cycles of 95°C for 15 s and 60°C for 60 s. The results for each gene were normalized by 18S rRNA levels measured in parallel in each sample.

### 4.6. Measurement of cytokine release

Concentrations of IL10, TNFα, IL6 and IL1β in the supernatant of microglial culture were measured using the ELISA kit (R&D systems) according to the manufacturer’s instructions. For the IL10 neutralization experiment, neutralizing antibody against IL10 (αIL10 rat IgG) or control antibody (control rat IgG) was added with other chemicals simultaneously.

### 4.7. MTT assay

Cell viability was assessed by a 3-(4,5-dimethly-2-thiazolyl)-2,5-diphenyltetrazolium bromide (MTT) assay, according to the manufacturer’s protocol (Nacalai Tesque). Microglia plated in 48-well plate were treated with drugs for 72 h. MTT (0.05 mg/mL) was then applied to the cultured cells for 2 h, followed by solubilization with dimethyl sulfoxide, and the absorbance at 570 nm was measured. Cell viability was expressed as a ratio to the control.

### 4.8. Statistical Analysis

All data were expressed as means ± SEM. Statistical analyses were performed by Student’s t-test or one-way ANOVA followed by Tukey’s test using GraphPad Prism 8. In all cases, differences of P < 0.05 were considered statistically significant.

## 5. Conclusions

In the present study, TRPV4 activation results in a shift in microglial activation state to IL10^high^ CD86^+^ leading to the enhancement of LPS-induced IL10 production in microglia. Our work reveals an important role of TRPV4-dependent IL10 production in the inhibition of pro-inflammatory cytokine production. These findings provide further insight into the relationship between microglia and cytokines.

## AUTHOR CONTRIBUTION

K.T. E.T. and H.S. conceptualized and designed the project; K.T. E.T. wrote the manuscript with the help of H.S.; K.T. E.T. S.Ku. and T.K. performed experiments; K.T.

E.T. and H.S. analyzed data; K.S. provided TRPV4-KO mice; H.S. supervised the project with help of K.N., S.Ka., and K.S.

## DATA AVAILABILITY STATEMENT

The data that support the findings of this study are available from the corresponding author upon reasonable request.

## ACKNOWLEDGMENTS

This work was supported by Grants-in-Aid for Scientific Research (KAKENHI) from MEXT/JSPS (to H.S., JP19H03377, JP23H02639, and JP23K27330), the Takeda Science Foundation (to H.S.), and the Uehara Memorial Foundation (to H.S.).

## References

1. Salter, M.W.; Stevens, B. Microglia emerge as central players in brain disease. Nat Med 2017, 23, 1018–1027, doi:10.1038/nm.4397.

2. Wang, W.Y.; Tan, M.S.; Yu, J.T.; Tan, L. Role of pro-inflammatory cytokines released from microglia in Alzheimer’s disease. Ann Transl Med 2015, 3, 136, doi:10.3978/j.issn.2305-5839.2015.03.49.

3. Colonna, M.; Butovsky, O. Microglia Function in the Central Nervous System During Health and Neurodegeneration. Annu Rev Immunol 2017, 35, 441–468, doi:10.1146/annurev-immunol-051116-052358.

4. Block, M.L.; Zecca, L.; Hong, J.S. Microglia-mediated neurotoxicity: uncovering the molecular mechanisms. Nat Rev Neurosci 2007, 8, 57–69, doi:10.1038/nrn2038.

5. Tang, Y.; Le, W. Differential Roles of M1 and M2 Microglia in Neurodegenerative Diseases. Mol Neurobiol 2016, 53, 1181–1194, doi:10.1007/s12035-014-9070-5.

6. Park, K.W.; Lee, H.G.; Jin, B.K.; Lee, Y.B. Interleukin-10 endogenously expressed in microglia prevents lipopolysaccharide-induced neurodegeneration in the rat cerebral cortex in vivo. Exp Mol Med 2007, 39, 812–819, doi:10.1038/emm.2007.88.

7. Chen, Z.; Trapp, B.D. Microglia and neuroprotection. J Neurochem 2016, 136 Suppl 1, 10–17, doi:10.1111/jnc.13062.

8. Kwilasz, A.J.; Grace, P.M.; Serbedzija, P.; Maier, S.F.; Watkins, L.R. The therapeutic potential of interleukin-10 in neuroimmune diseases. Neuropharmacology 2015, 96, 55–69, doi:10.1016/j.neuropharm.2014.10.020.

9. Anderson, W.D.; Greenhalgh, A.D.; Takwale, A.; David, S.; Vadigepalli, R. Novel Influences of IL-10 on CNS Inflammation Revealed by Integrated Analyses of Cytokine Networks and Microglial Morphology. Front Cell Neurosci 2017, 11, 233, doi:10.3389/fncel.2017.00233.

10. Arimoto, T.; Choi, D.Y.; Lu, X.; Liu, M.; Nguyen, X.V.; Zheng, N.; Stewart, C.A.; Kim, H.C.; Bing, G. Interleukin-10 protects against inflammation-mediated degeneration of dopaminergic neurons in substantia nigra. Neurobiol Aging 2007, 28, 894–906, doi:10.1016/j.neurobiolaging.2006.04.011.

11. Schwenkgrub, J.; Joniec-Maciejak, I.; Sznejder-Pachołek, A.; Wawer, A.; Ciesielska, A.; Bankiewicz, K.; Członkowska, A.; Członkowski, A. Effect of human interleukin-10 on the expression of nitric oxide synthases in the MPTP-based model of Parkinson’s disease. Pharmacol Rep 2013, 65, 44–49, doi:10.1016/s1734-1140(13)70962-9.

12. Bido, S.; Nannoni, M.; Muggeo, S.; Gambarè, D.; Ruffini, G.; Bellini, E.; Passeri, L.; Iaia, S.; Luoni, M.; Provinciali, M.;, et al. Microglia-specific. Sci Transl Med 2024, 16, eadm8563, doi:10.1126/scitranslmed.adm8563.

13. Waubant, E.; Gee, L.; Bacchetti, P.; Sloan, R.; Cotleur, A.; Rudick, R.; Goodkin, D. Relationship between serum levels of IL-10, MRI activity and interferon beta-1a therapy in patients with relapsing remitting MS. J Neuroimmunol 2001, 112, 139–145, doi:10.1016/s0165-5728(00)00355-6.

14. Bettelli, E.; Das, M.P.; Howard, E.D.; Weiner, H.L.; Sobel, R.A.; Kuchroo, V.K. IL-10 is critical in the regulation of autoimmune encephalomyelitis as demonstrated by studies of IL-10- and IL-4-deficient and transgenic mice. J Immunol 1998, 161, 3299–3306.

15. O’Neill, E.J.; Day, M.J.; Wraith, D.C. IL-10 is essential for disease protection following intranasal peptide administration in the C57BL/6 model of EAE. J Neuroimmunol 2006, 178, 1–8, doi:10.1016/j.jneuroim.2006.05.030.

16. Backonja, M.M.; Coe, C.L.; Muller, D.A.; Schell, K. Altered cytokine levels in the blood and cerebrospinal fluid of chronic pain patients. J Neuroimmunol 2008, 195, 157–163, doi:10.1016/j.jneuroim.2008.01.005.

17. Milligan, E.D.; Langer, S.J.; Sloane, E.M.; He, L.; Wieseler-Frank, J.; O’Connor, K.; Martin, D.; Forsayeth, J.R.; Maier, S.F.; Johnson, K.;, et al. Controlling pathological pain by adenovirally driven spinal production of the anti-inflammatory cytokine, interleukin-10. Eur J Neurosci 2005, 21, 2136–2148, doi:10.1111/j.1460-9568.2005.04057.x.

18. White, J.P.; Cibelli, M.; Urban, L.; Nilius, B.; McGeown, J.G.; Nagy, I. TRPV4: Molecular Conductor of a Diverse Orchestra. Physiol Rev 2016, 96, 911–973, doi:10.1152/physrev.00016.2015.

19. Zhang, F.; Mehta, H.; Choudhary, H.H.; Islam, R.; Hanafy, K.A. TRPV4 Channel in Neurological Disease: from Molecular Mechanisms to Therapeutic Potential. Mol Neurobiol 2025, 62, 3877–3891, doi:10.1007/s12035-024-04518-5.

20. Konno, M.; Shirakawa, H.; Iida, S.; Sakimoto, S.; Matsutani, I.; Miyake, T.; Kageyama, K.; Nakagawa, T.; Shibasaki, K.; Kaneko, S. Stimulation of transient receptor potential vanilloid 4 channel suppresses abnormal activation of microglia induced by lipopolysaccharide. Glia 2012, 60, 761–770, doi:10.1002/glia.22306.

21. Miyake, T.; Shirakawa, H.; Kusano, A.; Sakimoto, S.; Konno, M.; Nakagawa, T.; Mori, Y.; Kaneko, S. TRPM2 contributes to LPS/IFNγ-induced production of nitric oxide via the p38/JNK pathway in microglia. Biochem Biophys Res Commun 2014, 444, 212–217, doi:10.1016/j.bbrc.2014.01.022.

22. Kim, S.R.; Kim, S.U.; Oh, U.; Jin, B.K. Transient receptor potential vanilloid subtype 1 mediates microglial cell death in vivo and in vitro via Ca^2+^-mediated mitochondrial damage and cytochrome c release. J Immunol 2006, 177, 4322–4329, doi:10.4049/jimmunol.177.7.4322.

23. Nevelchuk, S.; Brawek, B.; Schwarz, N.; Valiente-Gabioud, A.; Wuttke, T.V.; Kovalchuk, Y.; Koch, H.; Höllig, A.; Steiner, F.; Figarella, K.;, et al. Morphotype-specific calcium signaling in human microglia. J Neuroinflammation 2024, 21, 175, doi:10.1186/s12974-024-03169-6.

24. Mosser, D.M.; Edwards, J.P. Exploring the full spectrum of macrophage activation. Nat Rev Immunol 2008, 8, 958–969, doi:10.1038/nri2448.

25. Mantovani, A.; Sica, A.; Sozzani, S.; Allavena, P.; Vecchi, A.; Locati, M. The chemokine system in diverse forms of macrophage activation and polarization. Trends Immunol 2004, 25, 677–686, doi:10.1016/j.it.2004.09.015.

26. Chhor, V.; Le Charpentier, T.; Lebon, S.; Oré, M.V.; Celador, I.L.; Josserand, J.; Degos, V.; Jacotot, E.; Hagberg, H.; Sävman, K.;, et al. Characterization of phenotype markers and neuronotoxic potential of polarised primary microglia in vitro. Brain Behav Immun 2013, 32, 70–85, doi:10.1016/j.bbi.2013.02.005.

27. MacKenzie, K.F.; Clark, K.; Naqvi, S.; McGuire, V.A.; Nöehren, G.; Kristariyanto, Y.; van den Bosch, M.; Mudaliar, M.; McCarthy, P.C.; Pattison, M.J.;, et al. PGE(2) induces macrophage IL-10 production and a regulatory-like phenotype via a protein kinase A-SIK-CRTC3 pathway. J Immunol 2013, 190, 565–577, doi:10.4049/jimmunol.1202462.

28. Edwards, J.P.; Zhang, X.; Frauwirth, K.A.; Mosser, D.M. Biochemical and functional characterization of three activated macrophage populations. J Leukoc Biol 2006, 80, 1298–1307, doi:10.1189/jlb.0406249.

29. Shibasaki, K.; Suzuki, M.; Mizuno, A.; Tominaga, M. Effects of body temperature on neural activity in the hippocampus: regulation of resting membrane potentials by transient receptor potential vanilloid 4. J Neurosci 2007, 27, 1566–1575, doi:10.1523/JNEUROSCI.4284-06.2007.

30. Alpizar, Y.A.; Boonen, B.; Sanchez, A.; Jung, C.; López-Requena, A.; Naert, R.; Steelant, B.; Luyts, K.; Plata, C.; De Vooght, V.;, et al. TRPV4 activation triggers protective responses to bacterial lipopolysaccharides in airway epithelial cells. Nat Commun 2017, 8, 1059, doi:10.1038/s41467-017-01201-3.

31. Shibasaki, K.; Ikenaka, K.; Tamalu, F.; Tominaga, M.; Ishizaki, Y. A novel subtype of astrocytes expressing TRPV4 (transient receptor potential vanilloid 4) regulates neuronal excitability via release of gliotransmitters. J Biol Chem 2014, 289, 14470–14480, doi:10.1074/jbc.M114.557132.

32. Watanabe, H.; Vriens, J.; Prenen, J.; Droogmans, G.; Voets, T.; Nilius, B. Anandamide and arachidonic acid use epoxyeicosatrienoic acids to activate TRPV4 channels. Nature 2003, 424, 434–438, doi:10.1038/nature01807.

33. Mori, M.; Aihara, M.; Kume, K.; Hamanoue, M.; Kohsaka, S.; Shimizu, T. Predominant expression of platelet-activating factor receptor in the rat brain microglia. J Neurosci 1996, 16, 3590–3600, doi:10.1523/JNEUROSCI.16-11-03590.1996.

34. Lin, D.; Gold, A.; Kaye, S.; Atkinson, J.R.; Tol, M.; Sas, A.; Segal, B.; Tontonoz, P.; Zhu, J.; Gao, J. Arachidonic Acid Mobilization and Peroxidation Promote Microglial Dysfunction in Aβ Pathology. J Neurosci 2024, 44, doi:10.1523/JNEUROSCI.0202-24.2024.

35. Rőszer, T. Understanding the Mysterious M2 Macrophage through Activation Markers and Effector Mechanisms. Mediators Inflamm 2015, 2015, 816460, doi:10.1155/2015/816460.

36. Walker, D.G.; Lue, L.F. Immune phenotypes of microglia in human neurodegenerative disease: challenges to detecting microglial polarization in human brains. Alzheimers Res Ther 2015, 7, 56, doi:10.1186/s13195-015-0139-9.

37. Franco, R.; Fernández-Suárez, D. Alternatively activated microglia and macrophages in the central nervous system. Prog Neurobiol 2015, 131, 65–86, doi:10.1016/j.pneurobio.2015.05.003.

38. Zhou, Z.; Peng, X.; Insolera, R.; Fink, D.J.; Mata, M. IL-10 promotes neuronal survival following spinal cord injury. Exp Neurol 2009, 220, 183–190, doi:10.1016/j.expneurol.2009.08.018.

39. Molina-Holgado, E.; Vela, J.M.; Arévalo-Martín, A.; Guaza, C. LPS/IFN-gamma cytotoxicity in oligodendroglial cells: role of nitric oxide and protection by the anti-inflammatory cytokine IL-10. Eur J Neurosci 2001, 13, 493–502, doi:10.1046/j.0953-816x.2000.01412.x.

40. Norden, D.M.; Fenn, A.M.; Dugan, A.; Godbout, J.P. TGFβ produced by IL-10 redirected astrocytes attenuates microglial activation. Glia 2014, 62, 881–895, doi:10.1002/glia.22647.

41. Gravel, M.; Béland, L.C.; Soucy, G.; Abdelhamid, E.; Rahimian, R.; Gravel, C.; Kriz, J. IL-10 Controls Early Microglial Phenotypes and Disease Onset in ALS Caused by Misfolded Superoxide Dismutase 1. J Neurosci 2016, 36, 1031–1048, doi:10.1523/JNEUROSCI.0854-15.2016.

42. Weston, L.L.; Jiang, S.; Chisholm, D.; Jantzie, L.L.; Bhaskar, K. Interleukin-10 deficiency exacerbates inflammation-induced tau pathology. J Neuroinflammation 2021, 18, 161, doi:10.1186/s12974-021-02211-1.

43. Hu, X.; Du, L.; Liu, S.; Lan, Z.; Zang, K.; Feng, J.; Zhao, Y.; Yang, X.; Xie, Z.; Wang, P.L.;, et al. A TRPV4-dependent neuroimmune axis in the spinal cord promotes neuropathic pain. J Clin Invest 2023, 133, doi:10.1172/JCI161507.

